# BRD4 expression in microglia supports recruitment of T cells into the CNS and exacerbates EAE

**DOI:** 10.1101/2024.09.13.612948

**Authors:** Anup Dey, Matthew Butcher, Anne Gegonne, Dinah S. Singer, Jinfang Zhu, Keiko Ozato

## Abstract

In EAE, a mouse model of multiple sclerosis, immunization with MOG autoantigen results in the generation of Th1/Th17 T cells in the periphery. MOG-specific T cells then invade into the central nervous system (CNS), resulting in neuronal demyelination. Microglia, innate immune cells in the CNS are known to regulate various neuronal diseases. However, the role of microglia in EAE has remained elusive. BRD4 is a BET protein expressed in microglia, whether BRD4 in microglia contributes to EAE has not been determined. We show that EAE pathology was markedly reduced with microglia-specific Brd4 conditional knockout (cKO). In these mice, microglia- T cell interactions were greatly reduced, leading to the lack of T cell reactivation. Microglia specific transcriptome data showed downregulation of genes required for interaction with and reactivation of T cells in Brd4 cKO samples. In summary, BRD4 plays a critical role in regulating microglia function in normal and EAE CNS.

**Summary:** This study demonstrates that in a EAE model, microglia-specific Brd4 conditional knockout mice were defective in expressing genes required for microglia- T cells interaction and those involved in neuroinflammation, and demyelination resulting in fewer CNS T cell invasion and display marked reduction in EAE pathology.

## Introduction

Multiple sclerosis (MS) is a neuroinflammatory disease of brain and spinal cord, caused by the infiltration of peripheral immune cells to the CNS from across impaired blood-brain barrier resulting in widespread demyelination of myelin sheath and damage to the neuroaxonal fibers (Lassmann, 2022). In experimental autoimmune encephalomyelitis (EAE), an animal model for MS, myelin specific CD4 autoreactive T cells are activated in peripheral lymph nodes, migrate to CNS and get reactivated by antigen presenting cells in the CNS (Prinz and Priller, 2017). Evidence shows that microglia, innate immune cells in the CNS play a critical role in modulating the brain parenchymal microenvironment. At steady state, microglia display a ramified morphology and engage in immune surveillance to help maintain CNS homeostasis. These functions have been ascribed to a distinct set of transcripts in the CNS (Healy et al., 2022; Hickman et al., 2013). In the context of neuroinflammation, microglia proliferate and assume morphology changes, changing from a ramified morphology to a shorter branched morphology. Following these morphological changes, microglia relocate to the site of injury or inflammation (Butovsky and Weiner, 2018; Masuda et al., 2020). In the EAE model, T cell entry to the CNS occurs with the migration and accumulation of other lymphocytes, including macrophages, dendritic cells, and monocytes to the sites of demyelination. Microglia, other CNS associated macrophages, and invading myeloid cells all may contribute to the pathology of EAE (Goldmann et al., 2013; Yoshida et al., 2014). In some studies it has been postulated that invading monocyte derived macrophages play a central role in the immune response and in demyelination, while resident microglia have been proposed to be responsible for surveying damage, reducing inflammation and removing debris (Yamasaki et al., 2014; Zabala et al., 2018). More recently, however, interaction between microglia and regulatory T cells has been shown to be critical at the relapsing-remitting model for EAE (Haimon et al., 2022). Direct crosstalk between microglia and T cells have been suggested to be associated with neurodegeneration (Chen et al., 2023). Furthermore, depletion of microglia by the colony stimulatory factor 1 receptor inhibitor, PLX5622 has been shown to diminish T cell proliferation, reactivation and delay in EAE onset (Montilla et al., 2023). The pathology of EAE depends on autoreactive CD4+ T cells, particularly Th17 and Th1/Th17 T cells that produce IL17A, IFNγ, GMCSF and TNFα. BET bromodomain inhibitor has been shown to control Th17 differentiation by regulating Th17 cytokines IL17A, IL21 and GMCSF and inhibited EAE symptoms in mice (Mele et al., 2013). These results suggested that BET bromodomain proteins (BRD2, BRD3, BRD4 and BRDT) either collectively or individually, may contribute to T cell differentiation in the periphery following immunization with the antigen. Thus, studies with BET inhibitors have not yet fully clarified whether BRD4 is critical for eliciting EAE pathology. BRD4 is distinguished from other BET proteins for its C-terminal specific binding to ‘positive transcription elongation factor’ (pTEFb) that recruits additional coactivators resulting in RNA PolII phosphorylation and transcription elongation of many genes (Jang et al., 2005; Kanno et al., 2014; Yang et al., 2005). *Brd4* expression is mostly ubiquitous with abundant expression from early embryos to most adult tissue (Bachu et al., 2016; Dey et al., 2000; Nishiyama et al., 2006; Shang et al., 2004). BRD4 is a cell cycle regulating protein that plays critical role in cell proliferation by regulating cell cycle specific genes (Wu et al., 2024). In prior studies, we demonstrated that BRD4 is critically required for hematopoiesis from fetal liver and bone marrow and modulate inflammatory gene expression in peripheral macrophages (Dey et al., 2019). Here, we constructed (i) CD4+ T cell and (ii) microglia-specific conditional Brd4 knockout (cKO) mice to determine the role of BRD4 in T helper cell activation in the peripheral lymph nodes and in microglia in the CNS, respectively. We first show that mice with T cell specific cKO do not develop EAE symptoms due to defective T cell priming and effector differentiation. Second. We show that mice with microglia specific Brd4 cKO were likewise resistant in developing severe EAE symptoms. Brd4 cKO microglia did not express genes expressed in wild type EAE microglia. Remarkably, Brd4 cKO microglia did not interact with T cells invading into the CNS and failed to reactivate T cells. Brd4 cKO microglia were defective in expressing genes required for the interaction with T cells and those involved in neuroinflammation and demyelination. Together, this study demonstrates a critical role of microglia and the requirement of BRD4 for eliciting EAE pathology.

## Results

### BRD4 expression in CD4 T cells is required for initial activation and subsequent infiltration of myelin specific effector T cells to CNS

By testing small molecule inhibitors, BET proteins have been identified to be critical for T cell activation (Bandukwala et al., 2012; Mele et al., 2013). These inhibitors target bromodomains of all members of BET family proteins and function in a dose dependent manner. In the EAE model, it is well established that, in response to immunization with MOG antigen, naïve T cells differentiate to Th17 cells within draining lymph nodes and are critically involved in the pathogenesis of EAE once they migrate to the CNS (Stadhouders et al., 2018). Previous investigation showed that transient treatment of JQ1, a BET inhibitor that prevents chromatin occupancy of BET proteins, inhibits transcription of Th17 genes and ameliorates EAE severity (Mele et al., 2013). To study the contribution of BRD4 excluding other BET proteins, we constructed Brd4 conditional knockout mice (cKO) lacking Brd4 in T cells (CD4-Cre; Brd4^f/f^). To generate EAE mice, 10 to 12 weeks old mice were immunized using an emulsion of myelin oligodendrocyte glycoprotein_35-55_ (MOG) peptide with complete Freund’s adjuvant and were co-injected with Pertussis toxin as described in the materials and methods section and in Figure 1A. On day 5 following immunization, cytokine expression was monitored in CD4 cells from draining lymph nodes. Total CD4 cell numbers were similar among all conditions, while IL17A expressing cells were significantly lower in Brd4 cKO mice relative to WT mice (see Cd4Cre Brd4^+/+^ - Brd4 WT and Cd4Cre Brd4^f/f^ - Brd4 cKO, Figure 1B, lower panels). No changes were observed between the genotypes when IFNγ expressing Th1 cells were considered (Figure 1B). Cells expressing both IL17A and IFNγ were not detectable at this stage (Figure 1B) (Stadhouders et al., 2018). WT mice developed EAE as expected with the onset of clinical symptoms on day 10 post immunization and reached the peak of EAE symptoms on day 18. Whereas Brd4 cKO mice were resistant to EAE, and they did not develop EAE symptoms within the 25-day experimental period (Figure 1C).

**Figure 1.**
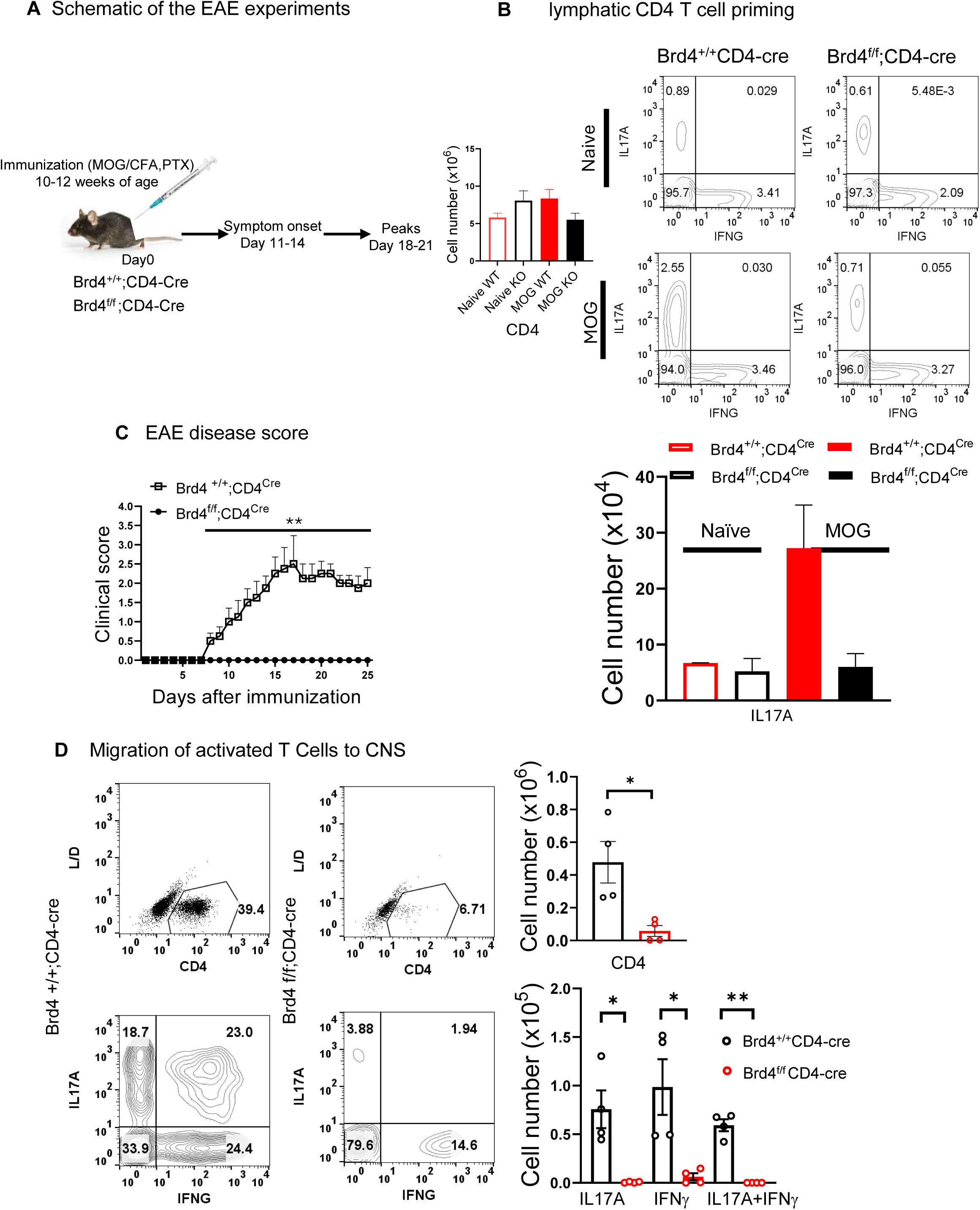
Brd4 deletion in CD4 T cells protects mice from EAE (A) Schematic showing EAE induction and its progression. Mice were immunized, and disease progression was monitored. (B) Cytokine expression 5-day post immunization from draining lymph nodes. T cell priming in MOG treated Brd4^+/+^; CD4^Cre/+^ (WT) mice was evidenced with significantly higher IL17 producing T cells, flow cytometric analysis (top), quantification of cell number below. Total number of T cells were similar (left). (C) Progression of EAE among Brd4^+/+^; CD4^Cre/+^ (WT) (n=4), while EAE symptoms were prevented in Brd4^f/f^ CD4^Cre/+^ (Brd4cKO mice) (n=4). Each data point is the average ± S.E.M., **p<0.001 (D) (Top left) Flow cytometric analysis of CNS invading CD4 T cells, Quantification of CD4 T cell number (right). (Bottom left) Flow cytometric analysis of corresponding T cells expressing IL17A, IL17A and IFNg and IFNg only, quantification on the right. Data presented as mean ± SD, **p<0.01, *p<0.05.

Examination of CNS cellularity reveals that in comparison to Brd4 cKO, there was a massive infiltration of CD4 T cells in WT (Figure 1D). Consequently, high number of CNS infiltrating CD4 T cells included those expressing IL17A alone, IFNγ alone or highly proinflammatory T helper cells coexpressing IL17A and IFNγ. These data show that Brd4 deletion in CD4 T cells prevented T cell activation and migration to the CNS thus preventing development of EAE. We conclude that BRD4 in Th cells promotes EAE pathogenesis.

### BRD4 deletion in microglia alone reduces EAE severity in mice

Naïve autoreactive T cells are activated during EAE by myelin oligodendrocyte glycoprotein_35-55_ (MOG) in draining lymph nodes triggering lymphoid cells and other myeloid cells to invade into the CNS through damaged blood-brain barrier. Upon entry to CNS, T cells come in contact with a milieu of antigen presenting cells and are activated further (Dong and Yong, 2019). While evidence indicates that microglia to contribute to EAE, this issue has remained elusive. Thus, to ask whether Brd4 in microglia affects EAE pathology: we constructed Brd4cKO mice using Cx3cr1^CreER^ Brd4^f/f^ strain. Brd4 was deleted upon tamoxifen injection (Figure 2A) (Goldmann et al., 2013). Vehicle (corn oil) injected Brd4^f/f^ Cx3Cr1-Cre mice were used as WT control. Microglia were identified by FACS analysis from spinal cord and brain (hereafter CNS) (Figure S1A). Myeloid cells were sorted as CD45^low^F4/80^+^TMEM119^+^ (microglia) and CD45^high^F4/80^+^TMEM119^-^ (invading myeloid cells). Efficient Brd4 deletion was confirmed, where in Brd4 cKO, CD45^low^ population (microglia), Brd4 transcript was not detectable, while robust Brd4 transcript was detected in WT (CD45^low^) and in CD45^high^ populations (Figure 2B). EAE clinical score for disease progression was monitored daily for 18 to 20 days. Graph in Figure 2C is an average of 3 independent sets of experiments, with 4 mice per group in each set. Clinical symptoms appeared in Brd4 WT mice around day12 and peaked by day 16-18. Brd4cKO mice exhibited milder symptoms during the period studied. This observation indicated that BRD4 function in microglia drives EAE pathogenesis. Demyelination in the white matter is one of the features of EAE pathogenesis. To determine the cause of milder EAE symptoms, we stained spinal cord sections from WT and Brd4 cKO mice with Luxol fast blue to determine the extent of myelin damage on the white matter. Widespread demyelination was evidenced throughout the length of WT spinal cord sections, while Brd4cKO spinal cord myelin layer was intact (Figure 2D). Shown on the right is the extent of demyelination, three data points each were taken from two mice. Noticeably, myelin damage in the WT accompanied massive influx of invading cells shown by the H & E stain, reduced cell migration was noticed in Brd4cKO throughout its length of the spinal cord (Figure S1B). Immunohistochemistry using Iba1 antibody detected a 2-5-fold increase in the number of microglia in the spinal cord of WT mice relative to Brd4 cKO mice (Figure 2E). This may be due to defective microglia proliferation in the early stages of EAE, given that BRD4 is known to promote proliferation (Plastini et al., 2020; Tan et al., 2022) (Wu et al., 2024). Morphologically, microglia from Brd4 cKO were highly ramified indicative of homeostatic surveillant type, while microglia from WT mice with EAE phenotype had numerous and shorter dendrites often seen in stress conditions (Figure 2E) (Vidal-Itriago et al., 2022).

**Figure 2.**
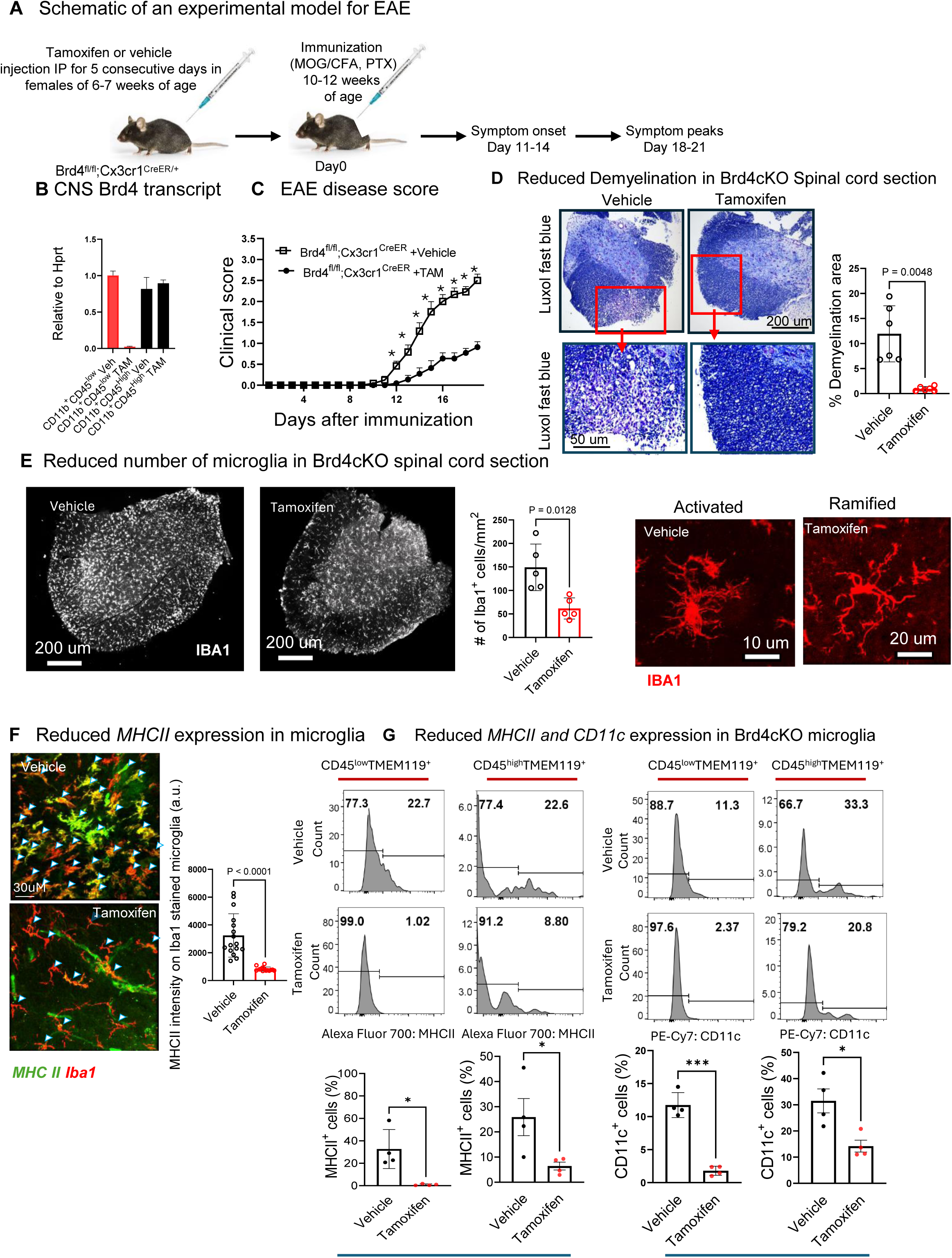
Effect of Brd4 deletion in microglia during EAE. (A) Schematic showing EAE induction and its progression. To delete Brd4, 6 to 7 weeks old Brd4^f/f^; Cx3cr1^Cre/+^ mice were treated with Tamoxifen for 5 consecutive days. 4 weeks post Tamoxifen or vehicle injection, mice were immunized, and disease progression was monitored. (B) qRT-PCR of Brd4 transcript extracted from microglia (CD45^low^CD11b^+^) and CNS invaded myeloid cells (CD45^High^CD11b^+^) isolated from WT (EAE) and Brd4cKO (healthy) mice respectively. Microglia and CNS invading myeloid cells were distinguished using surface markers CD45, CD11b, F4/80 and TMEM119 are presented in supplementary Figure 1A. (C) Following immunization EAE scores from Tamoxifen treated (Brd4cKO, reduced symptoms) and vehicle treated (WT, EAE) mice were recorded. Data represent average ± SEM of Tamoxifen (n=12) and vehicle (n=12) treated mice from 3 independent experiments *p<0.001. (D) Histology of spinal cord section using Luxol fast blue for assessing demyelination. Scale bars, 200um (top), 50um (below) showing intense demyelinated region on white matter in WT mice. Quantification of demyelination (right), data is average +/- S.D. from two independent experiments. (E) Immunohistochemistry of spinal cord section stained with Iba1 assessed microglia population (left). Quantification using ImageJ shows 2-fold increase in number in WT (middle) and change to reactive morphology (right). (F) Immunohistochemistry of spinal cord section showing colocalization of Iba1 stained microglia and MHCII. Intensity of MHCII was measured using ImageJ. MHCII staining was stronger in WT (vehicle). (Right) Strong MHCII expression is in WT (vehicle) (black symbol), while Brd4 cKO had weaker MHCII expression (Tamoxifen) (red symbol). (G) Flow cytometry plots (top) and corresponding quantification (bottom) of MHCII and CD11c expression from CD45^low/high^CD11b^+^TMEM119^+^ microglia. MHCII and CD11c expression were stronger in WT microglia (vehicle) on the left (black symbol) while were weaker in Brd4cKO (Tamoxifen) on the right (red symbol).

*MHCII* expression among various neural cells, have been implicated in many neurological diseases including MS (Harrington et al., 2023; Schetters et al., 2017). We addressed *MHCII* expression using immunohistochemistry and flow cytometry. Strong *MHCII* expression colocalizing I*ba1* staining were detected on WT microglia. *MHCII* expression was found to be much weaker in *Brd4* deficient microglia (Figure 2F). This observation was confirmed further by flow cytometry, where once again reduced *MHCII* expression was noted in Brd4 cKO microglia (Figure 2G, see CD45^low/high^ TMEM119^+^ cells). *CD11c* expressing microglia have been linked to aging and disease condition including MS and EAE (Benmamar-Badel et al., 2020). We observed reduced *CD11c* expression in *Brd4*cKO microglia (Figure 2G). Additionally, reduced expression of *CD80* costimulatory molecules was observed in Brd4 deleted microglia (Figure S1C). In contrast, expression of *MHCII, CD11c* and *CD80* remained high among invading myeloid cells (Figure S1D). These results showed that BRD4 in microglia, not infiltrating myeloid cells played a direct role in EAE development.

### Brd4 deletion in homeostatic Microglia

Brd4 is critically important in controlling differentiation and function of many immune cells, as it regulates transcription in a cell type specific manner (Dey et al., 2019; Gegonne et al., 2018; Yang et al., 2023). Given that Brd4 deletion in microglia caused profound impact on EAE pathology in mice, studying BRD4 regulated gene expression was of high interest. We performed whole genome transcriptome analyses for naïve microglia from WT and Brd4 cKO (Cx3cr1^CreER^;Brd4^f/f^) mice (Goldmann et al., 2013). CNS mononuclear cells were isolated 4 weeks after Tamoxifen treatment, microglia were identified as CD11b^+^CD45^low^F4/80^+^TMEM119^+^ and isolated by FACS sorting (Figure S2A). Purified single cell suspension were subjected for RNA-seq analysis. We used DESeq2 and used fold change > 2 and P<0.05 threshold to assess difference in the gene expression between WT and Brd4 cKO naïve microglia. With this criterion 720 genes were down regulated while 684 genes were up regulated in naïve Brd4 cKO microglia (Figure 3A). Gene ontology analysis using clusterProfiler software revealed down regulation of transcripts of various categories (Figure 3B) that includes genes related to mononuclear cell differentiation (*Dock10, Sox4, Bcl3, Il7r etc.*), mononuclear cell migration (Ccl7*, Ccl6, Ccr1, Ccl9, Ccl6, Trm2, Myo1g, Slamf1*) and positive regulation of leukocyte mediated immunity *(Nlrp3, Lag3, Il6, MR1)* (Figure 3C, D). These data indicated that BRD4 is required for establishing microglia response to external stimuli, cell migration and mediating immunity. However, it is important to note that transcription of homeostatic genes was unaltered between WT and Brd4 cKO (Figure S2B). A substantial number of genes were upregulated in Brd4 cKO microglia (684). GO analysis using clusterProfiler software identified these genes to: ATP metabolic process, generation of precursor metabolites and energy etc. (not shown)

**Figure 3.**
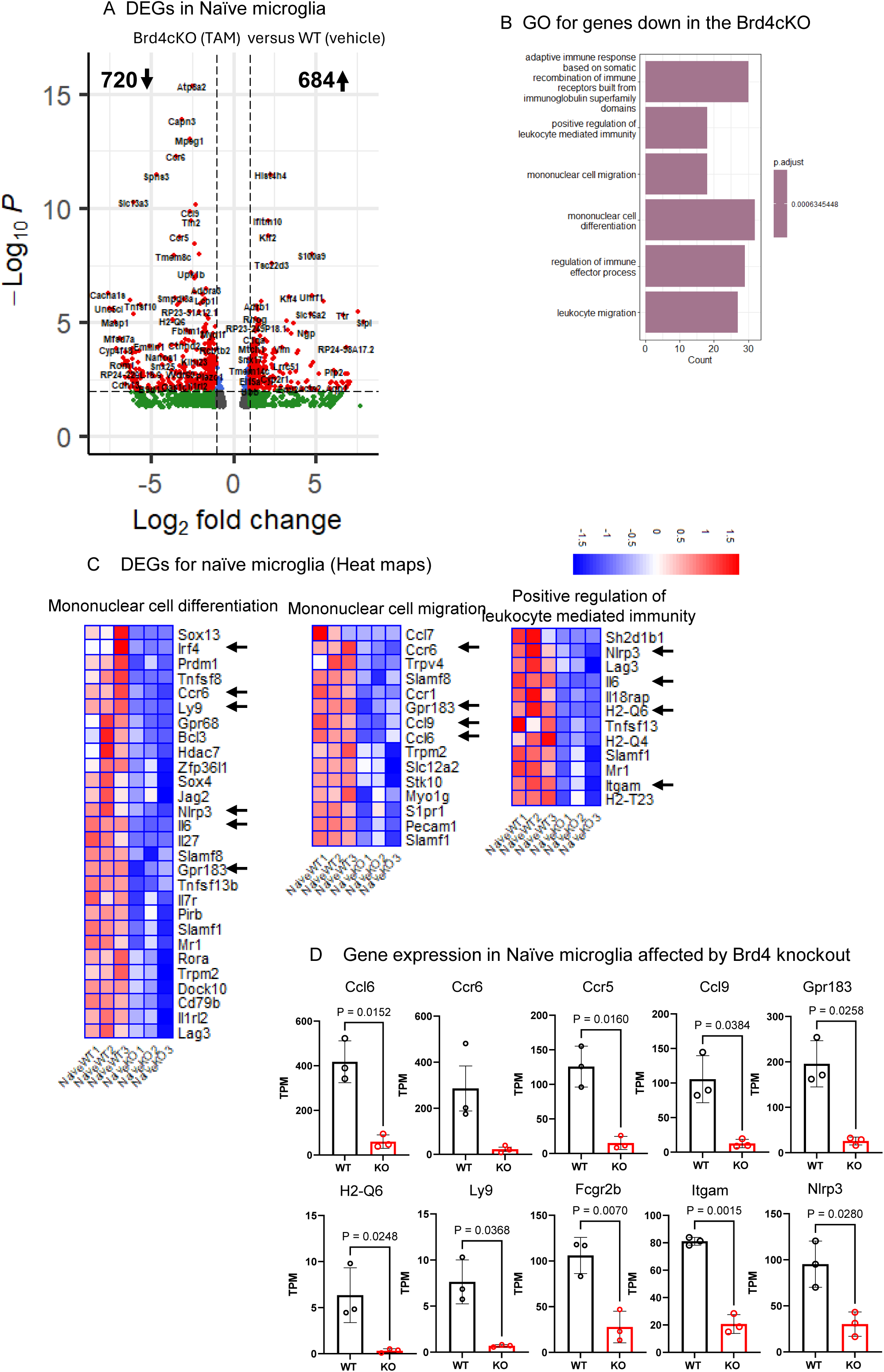
Transcriptome profiling of Brd4 deleted naïve microglia (A) Volcano plot comparing naïve microglia from Brd4cKO (Tamoxifen) and WT (vehicle) mice. Significant changes in expression (fold change >2, p value<0.05) are shown in red dots. (B) GO analysis of 720 downregulated genes in Brd4cKO (Tamoxifen) relative to WT (vehicle) CNS using clusterProfiler package in R. (C) Heat maps of genes from selected GO categories. (D) Quantification of relative TPM values of representative DEG’s selected from heat maps on C (arrows). Significance of differences were determined by unpaired t-test with Welch’s correction.

### Brd4 cKO microglia do not express genes induced after EAE

We next did RNA-seq analysis of WT and Brd4cKO microglia from EAE mice (18 days post MOG treatment) and estimated DEGs using the same criterion as above. Figure 4 and 5B show up- and down-genes from wild type EAE microglia, 343, 1015 respectively. Among down genes 177 were common to both naïve and MOG treated microglia (Figure S2C), but their relative expression was much higher in MOG treated microglia. Metascape software analysis revealed that these common down regulated genes are important for leucocyte activation, migration and immune response functions (Figure S2D). Microglia from MOG treated wild type CNS, had reduced expression of signature homeostatic markers such as, *P2ry12, Siglech, Gpr34, Hexb, Sall1, Tgfbr1, Mef2a, Mafb, Bin1, Smad3, (Butovsky and Weiner, 2018)* (cluster 1 Figure 4B, C, D), while they expressed at a normal level in Brd4cKO microglia. Concurrent with the loss of homeostatic microglia markers, there was a sharp increase in the expression of disease associated microglia genes (DAM), such as *Csf1, Il1b, Apoe, Cd74, Cst7, Axl, Lgals3* and *Fth1* in wild type, but not in *Brd4*cKO microglia (Figure 4B, C, D) Gene ontology enrichment analysis further revealed that down regulated transcripts overall were related to defense response, regulation of immune effector processes as well as antigen processing and presentation (Figure 5A).

**Figure 4.**
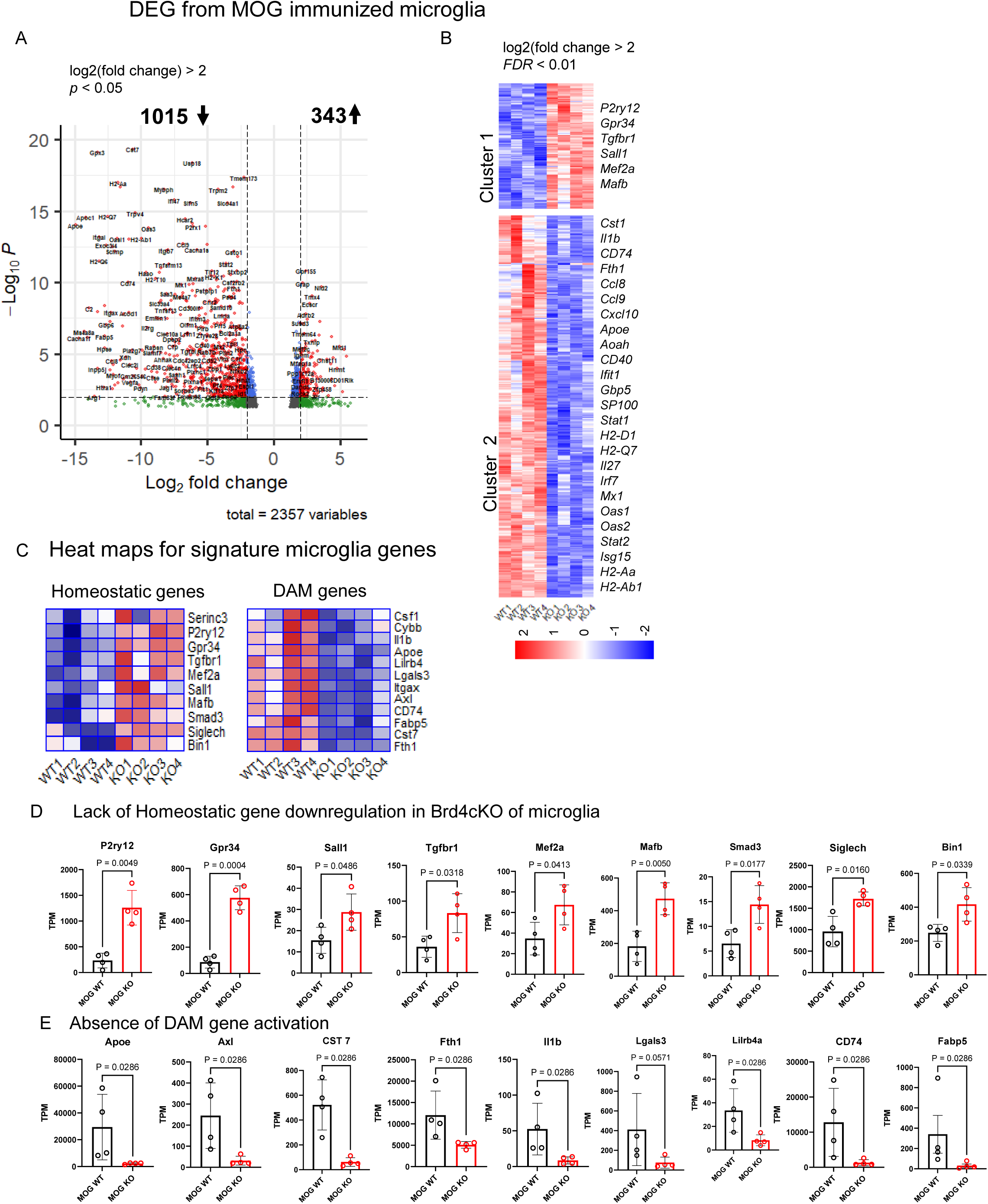
Unique transcriptional profile of Brd4cKO microglia during EAE (A) Volcano plot compares microglia gene expression from Brd4cKO (Tamoxifen) versus WT (vehicle) mice. Differential gene expression was determined using DESeq2 package, **s**ignificant changes in expression log2 (fold change) >2, p value<0.05) are shown in red dots. (B) Heat map derived from 867differentially expressed genes that were significantly altered in Brd4 KO microglia log2 (fold change) >2, FDR<0.01. (C) Hierarchical clustering distinguishing microglia identifying genes in homeostatic, and disease associated conditions. (D) Quantification of relative TPM values of representative DEG’s selected from heat maps on C. Significance of differences were determined by unpaired t-test with Welch’s correction.

**Figure 5.**
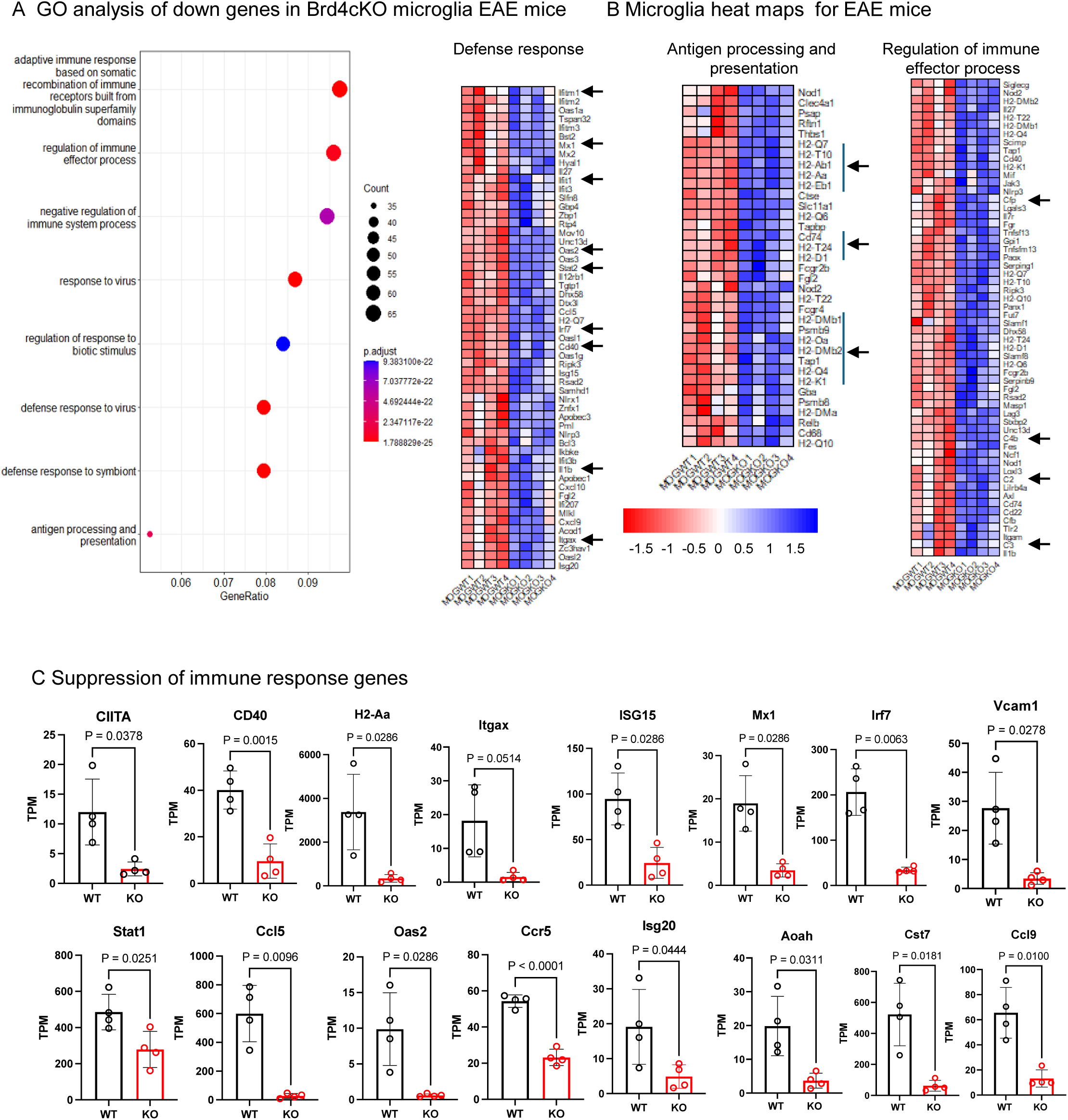
Contrasting microglial transcriptome from control and Brd4cKO Brd4^f/f^; Cx3cr1^Cre/+^ mice at the peak EAE. (A) GO analysis of 1015 genes down regulated in Brd4cKO microglia (from Figure 4), using clusterProfiler package. (B) Heat maps derived from genes in GO categories, such as defense response, antigen processing and presentation and regulation of immune effector process. (C) Quantification of relative TPM values of representative immune response genes from WT and Brd4 KO mice. Significance of differences were determined by unpaired t-test with Welch’s correction.

Function of microglia as an antigen presenting cell has been discussed in the past (Sosa et al., 2013; Wolf et al., 2018). In EAE for efficient adaptive immune response of T cells in the CNS, after its entry to CNS, T cells would require additional antigen presentation, cytokine signal from antigen presenting cells and costimulatory activity. Given that deletion of Brd4 from microglia rescued mice from severe disease, it may be hypothesized that BRD4cKO microglia are defective in antigen presentation. Consistent with this view *MhcII, Cd74, Ciita, Itgax (Cd11c)* were all down regulated in Brd4cKO microglia.

In addition, various chemokines such as, *Ccl5, Cxcl9 and Cxcl10,* genes in the complement system such as, *C2, C3, C4b* were downregulated. Likewose, interferon stimulated genes such as, *Mx1, Ifit1, Irf7, Oas2, IfitM1, Gbp6 and Stat1* were downregulated *in Brd4 cKO microglia* (Figure 5B and C). Various T cell costimulatory molecules expressed in microglia such as *CD40, CD80 and CD86* have been identified for T cell-microglia interaction. Upregulation of CD40 on microglia has been linked to T cell proliferation and infiltration of other leukocytes during EAE progression (Plastini et al., 2020). It was reported that in EAE and MS, Th cells express *CD40L* (Dong and Yong, 2019; Ponomarev et al., 2006). We found that *Cd40* gene expression was downregulated in Brd4cKO microglia, suggesting disruption in T cells-microglia interaction (Figure 5B, C). In other studies, with MS and EAE it was demonstrated that interaction of activated T cells and microglia via VCAM1 was important for TNF production from microglia causing inflammation, infiltration of peripheral immune cells and demyelination (Chabot et al., 1997; Dong and Yong, 2019). We found that Vcam1 expression was down regulated in Brd4 cKO microglia (Figure 5C). Aggravated demyelination was reported in cuprizone mediated MS using Cst7 knockout mice (Ling et al., 2019; Masuda et al., 2020). Cst7 was found downregulated in Brd4 cKO microglia, suggesting reduced demyelination in Brd4 cKO mice (Figure 5C).

### *Brd4 cKO* microglia do not interact with CNS invading T cells

Our findings that genes critical for microglia- T cells interaction was downregulated in Brd4 cKO microglia, prompted us to examine T cell number in the EAE CNS. First, we checked whether Brd4 deletion in microglia altered activated T cells in the periphery by testing T cells from spleen. FACS analysis revealed that Brd4 deletion in microglia did not affect the number of T cells in spleen, nor did the expression of *IL17A, IFNγ, GMCSF and TNFα* was changed (Figure S2E). This observation indicated that depletion of Brd4 in microglia did not alter activated T cell population in periphery. Next, we examined T cell population in the CNS. For detecting invading T cells, spinal cord sections were stained with CD3 antibody. In WT samples we detected accumulation of CD3 positive T cells, surrounded by *Iba1* positive microglia. In contrast only a few CD3 positive T cells were detected in the CNS of Brd4 cKO spinal cord (Figure 6A). The number of T cells shown on the right was an average of 2 sections from 4 mice corroborated reduction of CNS T cells. Reduction of T cell population in CNS was confirmed further by flow cytometry (Figure 6B). Additional FACS analysis in Figure 6C showed that the percentage of IL17A/IFN producing cells was lower in Brd4cKO CNS. These results are consistent with the view that Brd4cKO microglia do not interact with invading T cells, thus fail to reactivate them, leading to reduced EAE phenotype. Similar observation of reduced T cell population in CNS was reported earlier where PLX5622 was treated to deplete microglia and meningeal macrophages in EAE context (Montilla et al., 2023).

**Figure 6.**
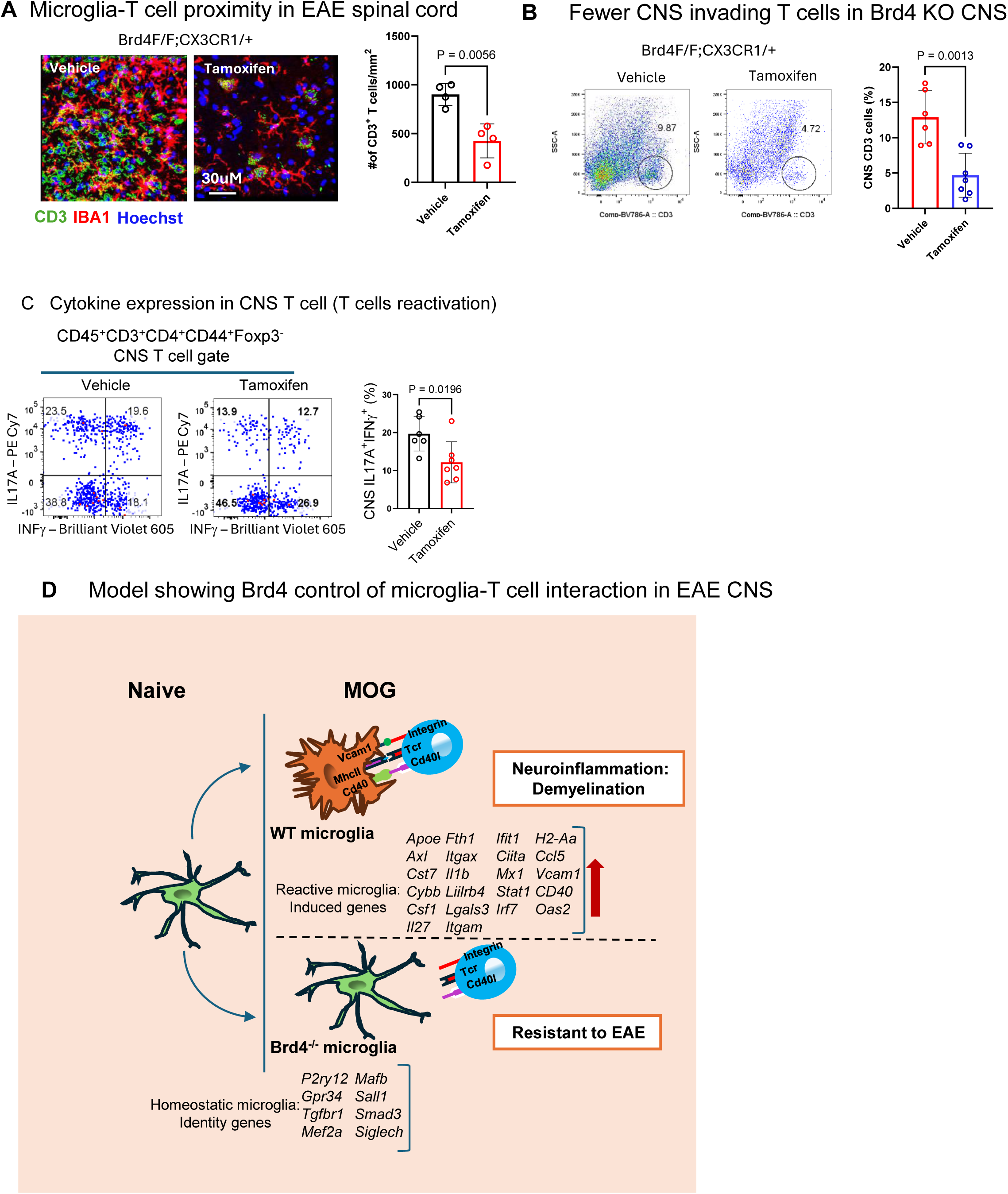
CNS infiltration of CD3 T cells from WT and Brd4cKO, Brd4^f/f^; Cx3cr1^Cre/+^ mice. (A) Immunohistochemistry of spinal cord section showing localization of Iba1 stained microglia and anti-CD3 labelled T cells. CD3 cell count using ImageJ shows increase in T cell number in WT sections (right). Significance of differences were determined by unpaired t-test with Welch’s correction. (B) Flow cytometry analysis showing reduced CD3 T cells in Brd4cKO CNS. Quantification of CD3 T cells (right), data was obtained from 6 WT and 7 Brd4cKO mice average +/- S.D. Significance of differences were determined by unpaired t-test with Welch’s correction. (C) Flow cytometry and quantification of CNS Th17A and IFNg cytokine expression (n>6). (D) Model for BRD4 control of microglia-T cell interaction in EAE CNS. In WT microglia, expression of Vcam1, MhcII, Cd40 promote interaction via integrin, TCR and CD40l on T cells. This leads to T cell proliferation and reactivation, microglia activation and upregulation of microglial proinflammatory and disease associated genes resulting in demyelination and axonal loss. Conversely, in Brd4 cKO microglia, homeostatic genes were not downregulated, DAM genes were not induced, but genes necessary for the interaction with T cells, neuroinflammation and demyelination are severely downregulated, resulting in reduced EAE pathology.

A diagram in Figure 6D depicts a model in which WT microglia interact with invading T cells, present MOG antigens, reactivate T cells and exacerbate EAE pathology. Whereas, Brd4cKO microglia do not interact with incoming T cells, nor reactivate them, thereby lessening EAE pathology (see Discussion for details).

## Discussion

In this study we investigated the function of BRD4 in TH1/Th17 effector T cells and microglia in the CNS during EAE. We first show that mice with Brd4cKO T cells do not develop EAE, indicating that BRD4 in T cells promotes EAE. Our results extend the previous report that BET inhibitors reverse EAE pathology and illustrate the importance of BRD4 among other BET proteins (Jahagirdar et al., 2017; Mele et al., 2013). Nevertheless, the central focus of this study was to explore the role of BRD4 in microglia in the context of EAE. We felt it is important to first clarify the contribution of microglia in EAE, since this issue has remained somewhat inconclusive. Namely, the role of microglia has been debated in the context of MHCII, as results could be attributed to infiltrated myeloid cells (Haimon et al., 2022; Wolf et al., 2018). We show that mice with Brd4cKO microglia, which carry WT peripheral myeloid cells, do not develop full EAE, rather show reduced demyelination, less neuroinflammation and reduced paralysis. Our data provide convincing evidence that microglia contribute to EAE progression. Our results are in line with the report that microglia ablation by PLX5622 inhibitor results in delayed onset and reduced EAE phenotype (Montilla et al., 2023).

Further, we show that BRD4 is a major factor that confers the functional activity of normal microglia. In addition, BRD4 helps drive EAE progression. Little, if any information has been available to the role of BRD4 in microglia so far. The present study provides convincing evidence that BRD4 is required for microglia function and transcriptome programs in health and disease. Besides microglia, BRD4 plays a critical role in gene expression and function of various other cell types. For example, BRD4 is required for the development of hematopoietic stem cells and modulates inflammatory responses in peripheral macrophages (Dey et al., 2019; Nicodeme et al., 2010). One of the most significant findings in this study was that Brd4 cKO microglia did not interact with invading T cells in the CNS. Earlier reports showed that in MS and EAE microglia and T cells were in proximity near the lesion, they interacted via cytokines and chemokines (Dong and Yong, 2019; Haimon et al., 2022) and the extent of damage was correlated by the number of MHCII carrying microglia and number of T cells (Androdias et al., 2010; Kuhlmann et al., 2002). Microglia ablation using PLX5622 inhibitor revealed a sharp decline in the number of infiltrated T cells, due to the decrease in T cell proliferation in the spinal cord resulting in delay in EAE onset (Heppner et al., 2005; Montilla et al., 2023). We found significantly fewer T cells in the CNS with Brd4 cKO microglia identified by immunohistochemistry and flow cytometry, indicating that BRD4 enables to maintain microglia-T cell interaction in the CNS. Transcriptome analysis provided mechanisms by which BRD4 controls microglia function under normal and EAE conditions. Brd4 cKO microglia did not express DAM genes that were expressed in WT microglia in EAE, nor did they downregulate homeostatic genes, indicating that BRD4 provides transcriptome programs specific for EAE. It has been shown that T cells infiltrate brain parenchyma via vascular cell adhesion protein (VCAM1) expressed on microglia (Androdias et al., 2010; Hauptmann et al., 2020). The reduction of invading T cells into the CNS observed with Brd4cKO microglia is likely explained at least in part by downregulation of Vcam1 expression.

Transcriptome data revealed that genes involved in antigen processing and presentation are downregulated in Brd4 cKO microglia, including chemokines, such as *Cxcl10, Ccl5*, antigen presentation such as, *Ciita, Cd74 and H2-2a*). Thus, Brd4 cKO microglia are defective in presenting MOG antigens to incoming T cells, therefore were unable to reactivate them. Furthermore, among other genes in the antigen presentation repertoire microglia expresses CD40, a costimulatory molecule that directly interact to T cells via CD40 ligand on T cells. It has been reported that this interaction is important for T cell expansion and continued infiltration of leukocytes to promote EAE disease progression (Plastini et al., 2020; Ponomarev et al., 2006). As CD40 expression was reduced in Brd4 cKO microglia, it would further undermine T cell-microglia interaction, resulting in inhibition of T cell expansion. Clinical symptoms in EAE accompanies demyelination of the CNS. Demyelination was reduced in Brd4 cKO mice. Significantly, we found demyelination related genes such as Cst7, Ccl6 and remyelination related genes such as MHC II and Cd74 were downregulated in Brd4 depleted microglia (Masuda et al., 2020). Further, inflammatory genes such as *Il1b, Ifit1, Mx1* were also downregulated in Brd4 cKO microglia, likely contributed reduced neuroinflammation (Figure 6D for a model).

In summary, our data reveal that BRD4 in microglia promotes EAE by directing multiple gene sets that collectively drive EAE progression.

## Materials and methods

### Mice

The generation of Brd4^f/f^ mice has been described previously by us (Dey et al., 2019). Brd4^f/f^ mice were backcrossed to C57BL/6 background and were crossed to CD4^Cre^ (Jackson Laboratory strain #022071) and Cx3cr1^CreER^ (Jackson laboratory strain #020940, kindly provided to us by Yosuke Mukoyama). For induction of Cre-mediated recombination 6 to 8 weeks old Brd4^f/f^; Cx3cr1^CreER/+^ females, were injected intra-peritoneally with 2mg Tamoxifen (Sigma), (20mg/ml in corn oil), for 5 consecutive days. Littermates with similar genotype were injected with 100ul corn oil and was used as WT control. All experiments were approved by National Institute of Child Health and Human development (Animal study Program #20-044, #23-044) and was carried out in accordance with National Institute of Health guidelines.

### EAE induction

Four weeks following Tamoxifen treatment, mice from both groups were immunized by subcutaneous injection with 200ug of MOG_35-55_, containing1-5mg heat killed Mycobacterium tuberculosis emulsified in Complete Freund’s adjuvant (EK-2110; Hooke Laboratories). Additionally, mice were injected with 100ng of pertussis toxin intraperitoneally and was repeated 24 hours later. Mice were scored daily based on detectable EAE symptoms; tip of the tail is limp - 0.5, limp tail - 1.0, limp tail hind leg inhibition – 1.5, hind limb paralysis - 2.0, front limb inhibition - 2.5, paralysis on both front and hind limb - 3.0.

### Immunohistochemistry

Animals were perfused transcardially with phosphate buffered saline (PBS) followed by 4% paraformaldehyde (PFA). Spinal cords were fixed further with PFA overnight and subsequently transferred to 30% sucrose in PBS. For immunofluorescence study 30um cryosections were blocked and permeabilized with 3% goat serum and 1% triton-X for 1 hour. For immunostaining sections were incubated overnight at 4^0^C with primary antibodies at a dilution of 1: 500 for rabbit anti Iba1 (Fujifilm), 1:500 for rat anti-CD3 (Abcam clone; CD3-12), 1:200 for rat anti-IA/IE (Biolegend clone; M5/114.15.2). For detection of these primary antibodies, as appropriate, sections were incubated for 2 hours at room temperature with Alexa fluor 488 and Alexa Fluor 633 conjugated goat secondary antibody (1:400, Invitrogen). Nuclei were detected by using Hoechst 33528 (1ug/ml, Sigma). Images were acquired using a Zeiss LSM800 confocal microscope to determine number and morphology of microglia and number of invading T cells in the white mater. Cell number was evaluated using ImageJ software.

### Histology

For histological analysis spinal cord were fixed with 4%PFA as described. Sections were embedded in paraffin before staining with Luxol fast blue and H&E to assess demyelination and neuronal invasion at the white mater. Histological images were obtained using a Zeiss Axiophot fluorescence microscope. Extent of demyelination were assessed using ImageJ software.

### Flow cytometry

For fluorescence activated cell sorting (FACS) of T cells in CD4 EAE model, cells were obtained from lymph nodes and stained with FITC CD4 (clone; GK1.5), PE/Cy7 IL17A (clone; TC11-18H10.1) and BV605 IFNγ (clone; XMG1.2). To analyze mononuclear cells from brain and spinal cord (CNS) cells were purified using a 70-30% neutral percoll gradient. Cells were then stained with Alexa Fluor 405 CD11b (clone: M1/70), APC/Cy7 CD45 (clone; 30-F11), BV605 F4/80 (clone: BM8), Alexa Fluor 488 TMEM119 (clone; 106-6), PE/Cy7 CD11c(clone; N418), Alexa Fluor 700 IA/IE (clone; M5/114), PE/Cy5 CD80 (clone;16-10A1) antibodies. Antibodies were obtained from Biolegend and was used in 1:100 dilution. Cells were analyzed using BD Fortessa or were FACS sorted using BD FACSAria. Data was analyzed using Flow Jo software, BD bioscience.

### RNA-Seq

For RNA-seq analysis 20-40,000 microglia cells were sorted directly in Trizol (Invitrogen), snaped freeze and stored at −80°C. RNA was isolated using Trizol-chloroform extraction. RNA-seq library was prepared using SmartSeq v4 Ultra Low input RNA Kit (Takara) and Illumina Nextera XT library preparation Kit. RNA-seq libraries were sequenced using NextSeq-500 system. 3 to 4 biological replicates were used in this study. Raw Fastq files were analyzed using NCBI pipeliner/4.0.2. For RNA-Seq analysis paired end reads were aligned to Mus musculus reference genome mm10, using splice-aware aligner STAR. DESeq2 software were used to determine differentially expressed genes with P-values and FDR. Transcript abundances were quantified using TPM.

### Statistical analysis

Data are presented here as mean ±SD with number of repeats showed in figures. All statistical analyses were performed using GraphPad Prism 9.0. Comparison between two groups were analyzed using unpaired t test with Welch’s correction. Statistical significance of EAE clinical score was determined by Mann Whitney nonparametric test. Statistically significant was considered when p-value was <0.05.

## Data availability

RNA-seq data are available at https://www.ncbi.nlm.nih.gov/geo/query/acc.cgi under accession number GSE275400.

## Acknowledgements

We like thank Yosuke Mukoyama (NHLBI) for kindly providing to us Cx3cr1^CreER^ mouse stain (Jackson laboratory strain #020940). We like to thank NIAID and NICHD FACS core facility for cell sorting. This research was supported by Intramural Research program of NICHD, NIH.

