## Supplemental figures for "BRD4 expression in microglia supports recruitment of T cells into the CNS and exacerbates EAE"

### Supplementary Figure 1

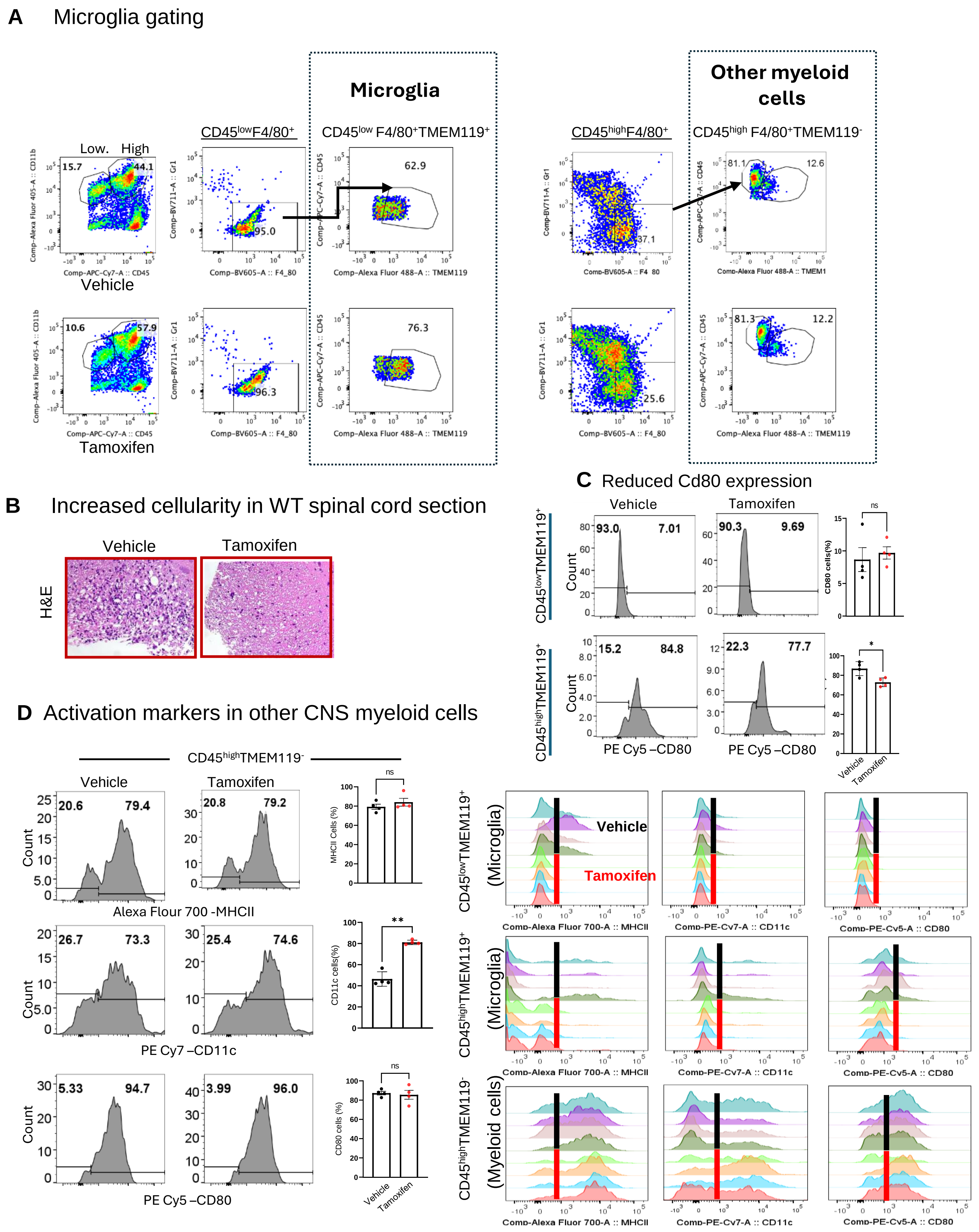

**Supplemental Figure 1** Gating strategy for ex-vivo analysis of microglia. (A) representative strategy for flow cytometry analysis and microglia sorting for RNA-seq. (B) Histological section of spinal cord stained with H&E. Influx of invading cells were detected in WT section shown as black dots, reduced in Brd4cKO section. (C) Flow cytometry plots and quantification of CD80 co stimulatory molecule showing elevated CD80 expression in $CD45^{high}CD11b^{+}TMEM119^{+}$ WT microglia but not in Brd4cKO microglia. (D) (Left) Flow cytometry quantification of MHCII, CD11c, CD80 from $CD45^{high}CD11b^{+}TMEM119^{-}$ myeloid cells. (Right) Flow cytometry quantification of MHCII, CD11c, CD80 from CD45 $^{low/high}CD11b^{+}TMEM119^{+}$ microglia and CD45 $^{high}CD11b^{+}TMEM119^{-}$ myeloid cells from CNS.

### Supplementary Figure 2

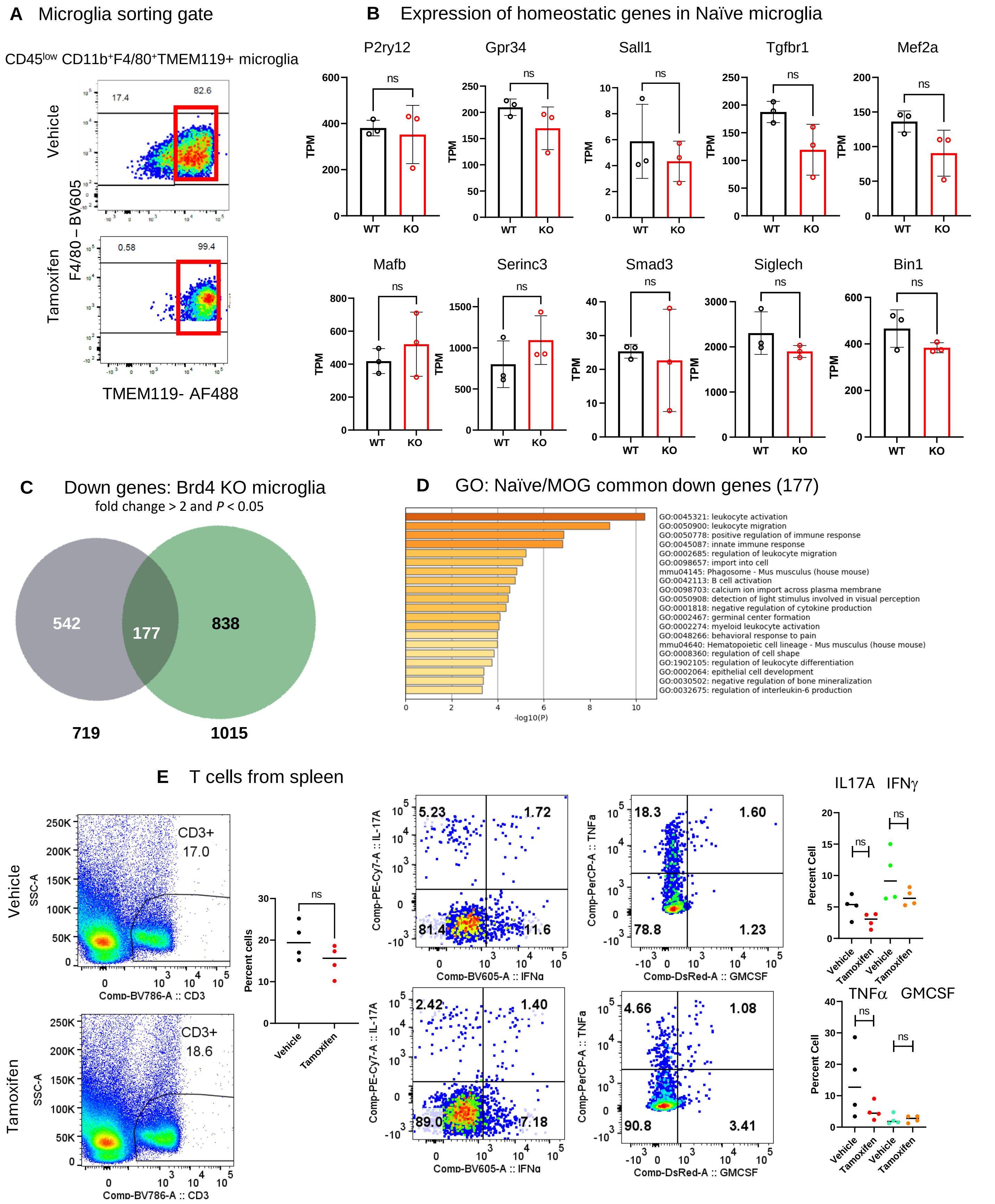

**Supplemental Figure 2** (A) Microglia sorting strategy (B) Quantification of relative TPM values of representative homeostatic genes from Naïve microglia. Significance of differences were determined by unpaired t-test with Welch's correction. (C) Venn diagram comparing down regulated genes from Naïve and MOG immunized microglia. (D) GO analysis of 177 down regulated genes common to naïve and MOG immunized microglia. (E) Flow cytometric analysis of peripheral T cells from spleen of MOG immunized mice. (right) Quantification of T cell percentages expressing IL17a, IFNg, TNFa and GMCSF are not statistically different among vehicle and Tamoxifen treated samples.
